# Real-time observation of light-controlled transcription in living cells

**DOI:** 10.1101/132050

**Authors:** Anne Rademacher, Fabian Erdel, Jorge Trojanowski, Karsten Rippe

**Affiliations:** German Cancer Research Center (DKFZ) and BioQuant, Research Group Genome Organization & Function, Im Neuenheimer Feld 280, 69120 Heidelberg, Germany

## Abstract

Gene expression is a tightly controlled process that is coordinated in space and time. To dissect its dynamic regulation with high temporal resolution, we introduce an optogenetic tool termed BLInCR (Blue Light-Induced Chromatin Recruitment) that combines rapid and reversible light-dependent recruitment of effector proteins with a real-time readout for transcription. We used BLInCR to control the activity of a reporter gene cluster in the human osteosarcoma cell line U2OS by reversibly recruiting the viral transactivator VP16. RNA production was detectable ∼2 minutes after VP16 recruitment and readily decreased when VP16 dissociated from the cluster in the absence of light. Quantitative assessment of the activation process revealed biphasic activation kinetics with a pronounced early phase in cells treated with the histone deacetylase inhibitor SAHA. Comparison with kinetic models for transcription activation suggests that the gene cluster undergoes a maturation process when activated. BLInCR will facilitate the study of transcription dynamics in living cells.

## Introduction

Tools that enable the accurate control, visualization and quantitation of the transcription process have driven recent progress in the field and are key to dissect its molecular underpinnings. Tracing RNA by live cell imaging has provided a wealth of information on gene regulation and RNA processing (Cho et al., 2016; Darzacq et al., 2009; Martin et al., 2013). In particular, a reporter gene array integrated into the human U2OS cell line has been used to investigate transcription activation and the associated changes in the chromatin environment (Janicki et al., 2004; Rafalska-Metcalf et al., 2010) as well as elongation and pausing by RNA polymerase II (Darzacq et al., 2007). However, elucidating the early steps of the activation process is difficult to accomplish with the existing techniques that rely on chemicals for transcription initiation and therefore depend on diffusion and uptake/release of the respective compounds. To control protein interactions with high temporal precision a variety of optogenetic methods have been adapted for use in mammalian cells (Tischer and Weiner, 2014). These include the CIBN-PHR system derived from the *Arabidopsis thaliana* proteins CIB1 and CRY2. Rapid binding of PHR-fused effector proteins to tethered CIBN can be induced by illumination with blue light (Kennedy et al., 2010). Different variations of the CIB1-CRY2 system have been exploited to recruit the viral transactivator VP64 to gene promoters marked by transcription activator-like effectors (TALEs) or dCas9 fusion constructs (Konermann et al., 2013; Polstein and Gersbach, 2015). However, these systems did not include live-cell readouts for transcriptional activity and were therefore not suited to study the kinetics of transcription activation with high time resolution. Here, we introduce a tool termed **B**lue **L**ight-**In**duced **C**hromatin **R**ecruitment (BLInCR), which combines rapid and reversible binding of effectors with a real-time transcription readout in living cells. We used BLInCR to dissect the transcription activation process at a gene cluster and to probe the persistence of its activated state.

## Results

### Implementation of a light-induced chromatin recruitment system

BLInCR is based on the PHR and CIBN domains of the *Arabidopsis thaliana* proteins CRY2 and CIB1 (Kennedy et al., 2010) that interact with each other when illuminated with blue light (Fig. 1A). Accordingly, CIBN fusion proteins that localize to nuclear structures or genomic loci of interest (‘localizers’) reversibly bind PHR fusion proteins (‘effectors’) upon blue light exposure. To test the versatility of the BLInCR system we triggered and visualized the targeting of fluorescently labeled effector proteins to different nuclear subcompartments in the human U2OS 2-6-3 cell line. It carries a stably integrated array of ∼200 reporter construct units with promoter-proximal repeats of the *tet*O bacterial operator sequence (Janicki et al., 2004). BLInCR robustly induced accumulation of PHR-mCherry as a mock effector at the reporter array, telomeres, nucleoli, PML bodies or the nuclear lamina (Fig. 1B, Supplementary Table S1). We set out to test whether BLInCR is compatible with the U2OS 2-6-3 reporter system (Fig. 2A) that allows for the detection of RNA transcripts at the reporter array via MS2 stem-loop sequences visualized by binding of fluorescently labeled MS2 coat proteins. The protein products encoded by this transcript contain a cyan fluorescent protein (CFP) domain and localize to peroxisomes. We transfected cells with CIBN-TetR and PHR-YFP-VP16, which contains the strong viral VP16 transcription activation domain, and illuminated them with blue light overnight. Both MS2-RNA accumulation at the reporter array and peroxisomal CFP expression were observed in almost all cells (RNA: 92%, CFP: 81%, *n* = 37; see Fig. 2B for a representative cell). RNA production at the array was confirmed by RNA FISH with a probe directed against the MS2 loop sequences of the transcript (Supplementary Fig. S1A,B).

**Figure 1.**
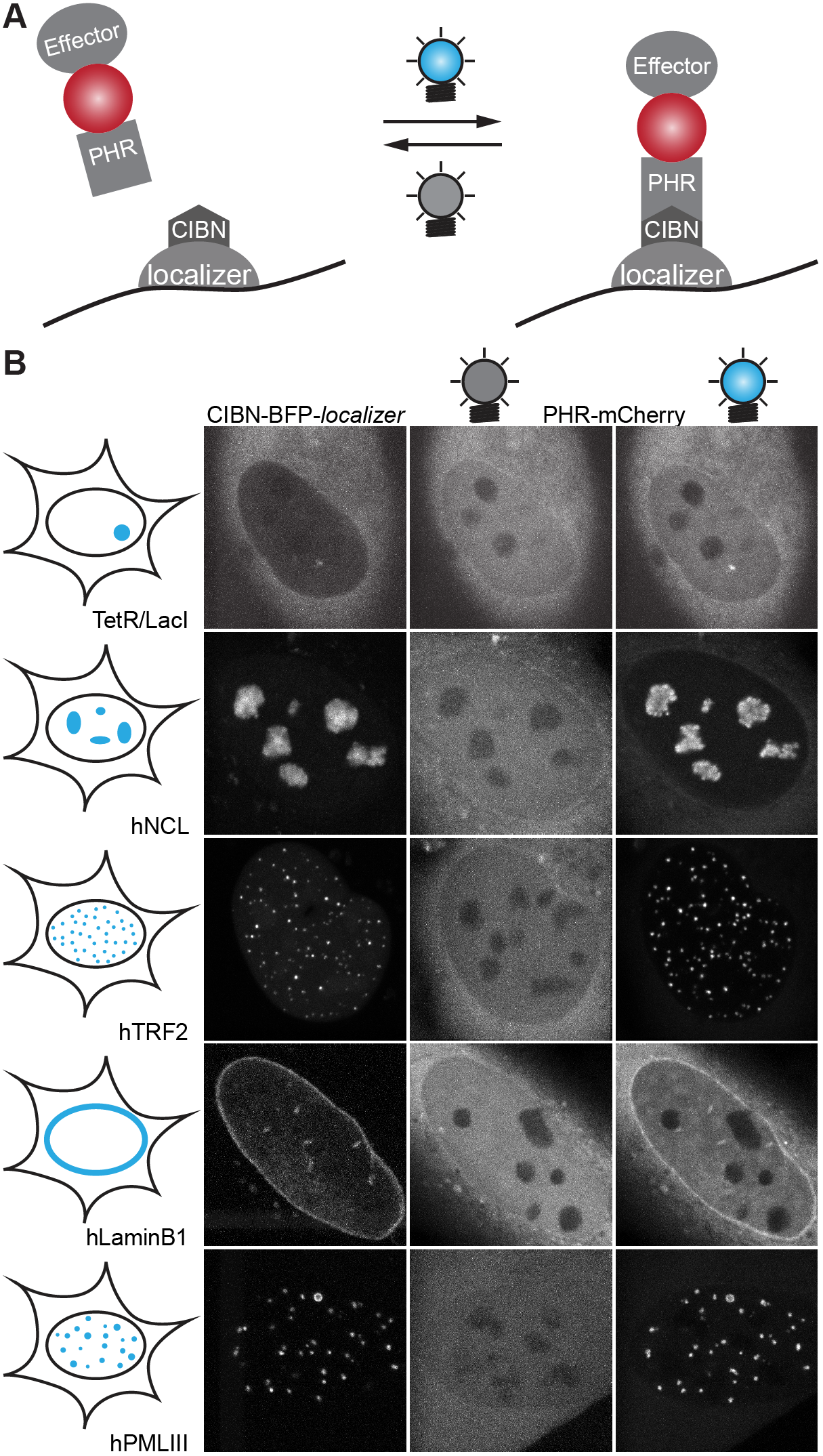
Blue Light-Induced Chromatin Recruitment (BLInCR). (**A**) BLInCR is based on the interaction of the protein domains CIBN and PHR upon illumination with blue light. It allows binding of PHR-tagged effectors to CIBN-marked sites. (**B**) BLInCR enables recruitment to different genomic loci: tetO arrays, nucleoli, telomeres, the nuclear lamina or PML bodies. PHR-mCherry (a mock effector) was homogeneously distributed throughout the cell before exposure (center) and relocated to sites marked by CIBN-BFP-tagged localizers (left) upon blue light exposure (right).

**Figure 2.**
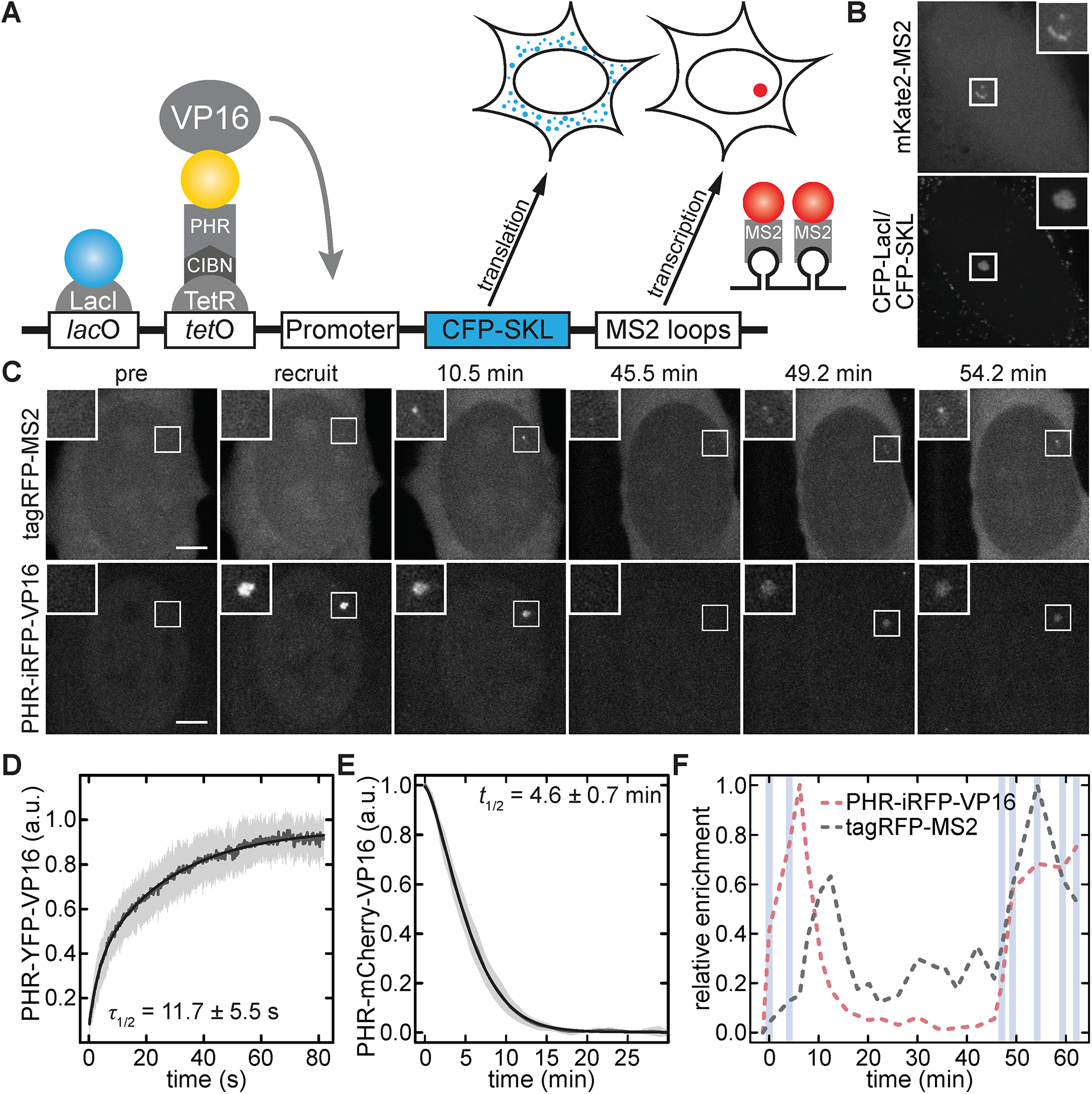
Dynamic transcription control by BLInCR. (**A**) Schematic representation of the reporter system used to detect transcription in real-time (Janicki et al., 2004). (**B**) Constitutive recruitment of the transcriptional activator VP16 induced production of RNA (top) and the encoded CFP-SKL protein (bottom). (**C**) Pulsed recruitment of VP16 (top) induced pulses of RNA accumulation visualized by labeled MS2 coat proteins (bottom). Scale bars: 5 μm. (**D**) Association kinetics of PHR-YFP-VP16 with CIBN-TetR-RFP tethered to the *tet*O reporter array. Averaged experimental data (dark gray lines), fits (black lines) and standard deviations (light gray areas) are shown (*n*=20). (**E**) Dissociation kinetics of the PHR-mCherry-VP16 and CIBN-TetR complex (*n*=13). Plot coloring as in panel **D**. (**F**) Quantification of the integrated signal at the array in panel **C**. Shaded areas represent blue light pulses.

### Kinetics of light-induced chromatin association and dissociation

To characterize the kinetics of light-induced association and subsequent dissociation of the VP16 effector protein at the reporter array, cells were transfected with a CIBN-TetR construct and a fluorescently labeled PHR-VP16 fusion. PHR-VP16 readily bound to the reporter array in the presence of blue light and dissociated in the dark (Fig. 2C, top), allowing for the implementation of tailored recruitment and activation patterns. To quantitate the association reaction, we first identified the array using RFP-tagged CIBN-TetR excited by green laser light, which does not induce optogenetic recruitment, and subsequently induced and recorded the accumulation of YFP-labeled PHR-VP16 at 200 ms time resolution with blue laser light (Supplementary Fig. S2A, **Movie 1**). The characteristic time to reach half-maximal levels amounted to 11.7 ± 5.5 s (Fig. 2D), which is about one and two orders of magnitude faster than doxycycline- and tamoxifen-induced recruitment, respectively (Normanno et al., 2015; Rafalska-Metcalf et al., 2010). YFP-labeled PHR fused to a nuclear localization sequence (NLS) or to the human histone acetyltransferase GCN5 accumulated with similar kinetics (Supplementary Fig. S2B, Supplementary Table S2). Induced accumulation at the array occurred fast with two characteristic rates (Supplementary Table S2), which might reflect the kinetics of the CIBN-PHR interaction and the previously described PHR oligomerization (Bugaj et al., 2013) that are both triggered by blue light. To quantitate the dissociation reaction, PHR-mCherry-VP16 was targeted to the reporter array in cells expressing CIBN-TetR by illumination with blue light for 38 s (Supplementary Fig. S2C). Subsequently, the loss of mCherry signal from the array was monitored over time, yielding a characteristic half-life of 4.6 ± 0.7 min (Fig. 2E), which was identical for PHR-mCherry-NLS (4.5 ± 0.5 min, Supplementary Fig. S2D, Supplementary Table S3). Notably, the effectors were completely dissociated from the array within 15-20 min.

### Persistence of transcriptional activation

To assess if transcriptional activity persisted after removal of the activating stimulus, we analyzed the RNA signal at the array (Fig. 2C, bottom) during and after VP16 recruitment. RNA production was readily detected upon VP16 arrival and decreased after VP16 removal (Fig. 2F). For the cell shown here, RNA production was more rapidly reactivated when inducing VP16 binding again after 40 minutes, which might be indicative of transcriptional memory. This effect could be due to a low level of sustained transcriptional activity, which persisted after the first activation and led to the retention of the transcription machinery, and/or due to changes in the chromatin landscape that predispose the locus to reactivation. These results show that BLInCR is a versatile tool to rapidly induce protein binding to nuclear structures and to tune its residence time on the timescale of minutes. It combines the capability of plant-based optogenetic systems for transcriptional control (Konermann et al., 2013; Motta-Mena et al., 2014; Niopek et al., 2014; Polstein and Gersbach, 2015) with the power of real-time microscopy readouts, e.g. (Janicki et al., 2004), making it possible to assess cellular function at subcellular resolution over time.

### Quantitation and modeling of transcription activation kinetics

Next, we used BLInCR to follow the transcription activation kinetics of the reporter array in U2OS 2-6-3 cells with high temporal resolution. Low RNA levels were already detectable after about two minutes (Fig. 3A), followed by a lag phase with little or no additional RNA accumulation (Fig. 3B, black line). After 20-30 minutes, a second phase of rapid accumulation of reporter RNA was observed, indicating that the activation process involves at least two distinct time scales. To assess the influence of activating chromatin marks on the activation kinetics we treated cells with the histone deacetylase inhibitor SAHA, which led to globally elevated histone acetylation levels (Fig. 3C). Local enrichment of histone acetylation has previously been shown to coincide with transcriptional activation of the reporter array (Rafalska-Metcalf et al., 2010) and was also observed when activating transcription using BLInCR overnight (Supplementary Fig. S3A-C). Cells that had been pretreated with SAHA exhibited a much more pronounced fast activation phase (Fig. 3B), indicating that preexisting histone acetylation facilitates the activation process. These results are consistent with the notion that some of the ∼200 copies of the reporter array are poised for immediate activation, and that this fraction can be increased by hyperacetylation of histones in SAHA-treated cells.

**Figure 3.**
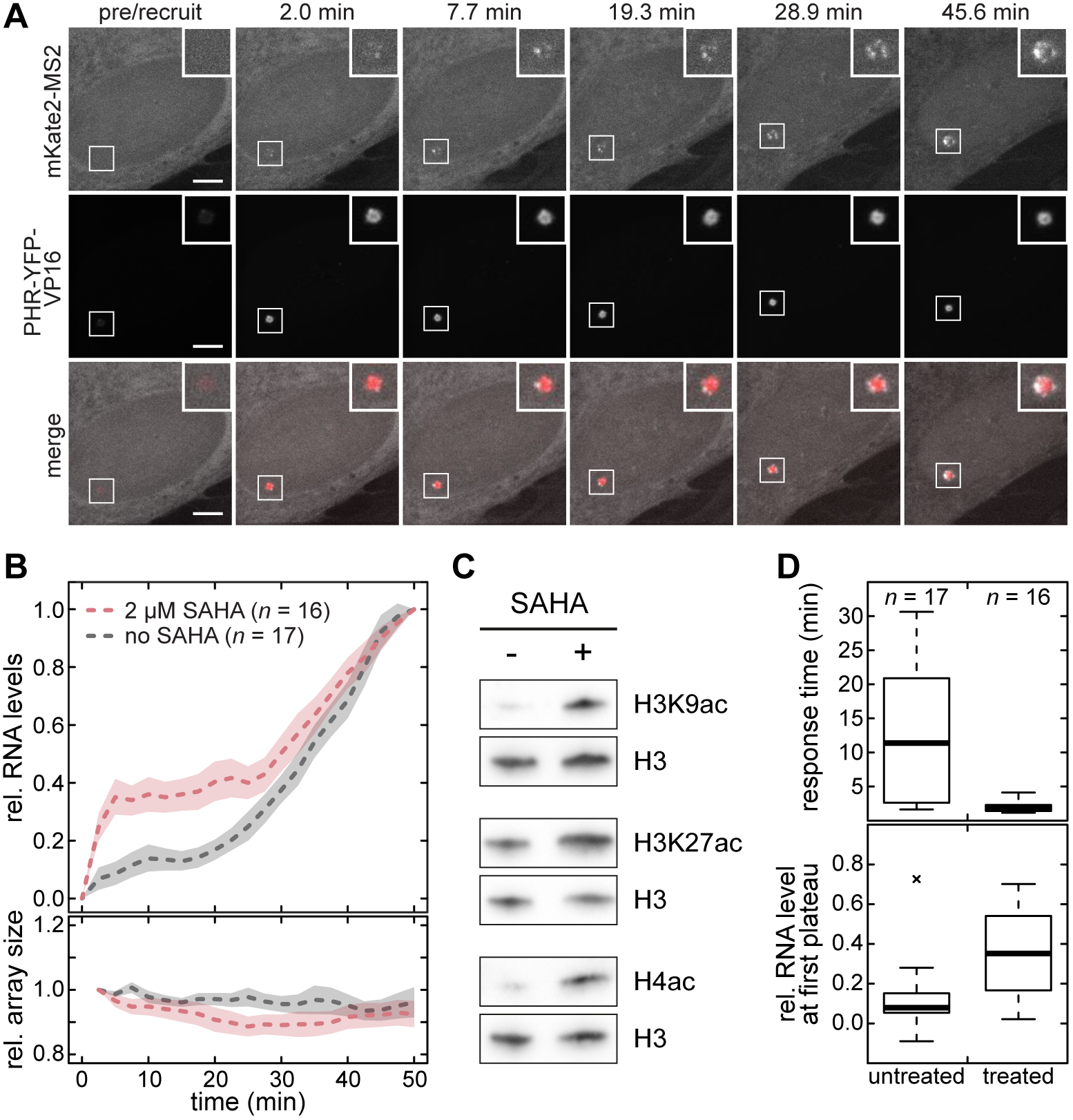
BLInCR resolves the transcription activation process with high resolution. (**A**) Time series of VP16-induced transcription. Top: RNA production visualized by labeled MS2 coat proteins. Center: PHR-YFP-VP16 was continuously present at the array during imaging. Bottom: Merged images. Scale bars: 5 μm. (**B**) Top: BLInCR-induced transcription occurred with an early and a late activation phase. The early response was more pronounced in cells pretreated with SAHA overnight. Depicted are experimental averages (dashed lines) and standard errors of the mean (shaded areas). Bottom: Relative size of the reporter array over time. (**C**) SAHA treatment at 2 μM concentration for 24 hours led to a reduction of preexisting histone acetylation marks. (**D**) Cell-to-cell heterogeneity of response times (top) and relative RNA levels (bottom) in treated and untreated cells. Response times correspond to the time points at which first transcripts were detectable. Relative RNA levels were measured at the point of inflection, i.e. at the first plateau between early and late transcription.

The produced reporter RNA was not homogenously distributed across the array (Fig. 3A, insets), suggesting that activated transcription sites might be clustered and co-regulated. The size of the reporter array remained unchanged within the first 50 minutes after PHR-YFP-VP16 recruitment (Fig. 3B, bottom). Thus, the global chromatin decompaction observed previously (Janicki et al., 2004; Rafalska-Metcalf et al., 2010) was not a prerequisite for PHR-YFP-VP16 mediated activation, but might rather occur downstream of transcription initiation (Supplementary Fig. S1C). The transcriptional response to VP16 recruitment was heterogeneous among cells. Without treatment, the response time, after which transcripts were first detectable, varied from less than two minutes to more than 30 minutes, whereas it was always less than 5 minutes in SAHA-treated cells (Fig. 3D, top). In addition, the extent of early transcription as estimated from the level of the first plateau at ∼9-12 min (see Fig. 3B) varied greatly for both treated and untreated cells (Fig. 3D, bottom) and heterogeneity was also observed among constitutively activated cells (Supplementary Fig. S1A,B). This heterogeneity might reflect differences in epigenetic promoter signals and other chromatin features.

To interrogate the underlying transcription activation mechanism, we analyzed the activation kinetics measured for treated and untreated cells (Fig. 3B) according to the theoretical framework described in the Methods section. In a comparison of different models, we found that a previously derived two-state model (Shahrezaei and Swain, 2008) did not fit the observed kinetics (Fig. 4A, left). This model was recently found to describe the reactivation kinetics of a single-copy reporter that was silenced by recruitment of a histone deacetylase beforehand (Bintu et al., 2016). In contrast, a model with the same number of parameters that included positive feedback and a fraction of promoters that acted as independent transcription units fitted the data well (Fig. 4A, center). Positive feedback could originate from spatial interactions among cooperatively transcribed reporter genes (Li et al., 2012; Papantonis and Cook, 2013) and might include the transcription-induced relocalization of promoters during the activation process (Therizols et al., 2014). A model that represents the activation process as a series of multiple sequential reaction steps with the same transition rate also yielded a good but slightly worse fit (Fig. 4A, right). Fit parameters for the different models are listed in Supplementary Table S4.

**Figure 4.**
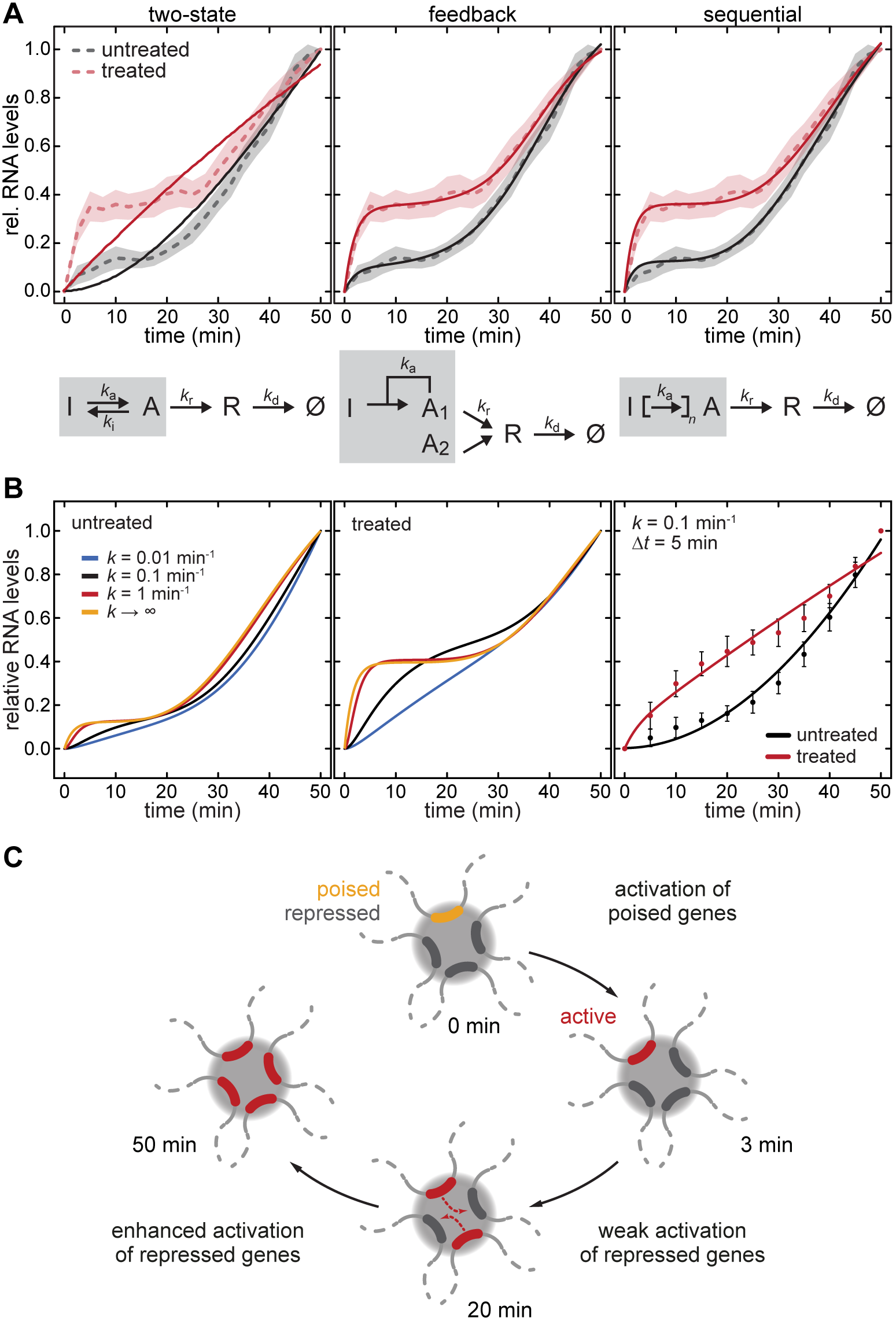
Kinetic model for transcription activation. (**A**) The experimentally determined activation kinetics were fitted with three different kinetic models (from left to right): a two-state model, a model including positive feedback, and a sequential activation model. The biphasic activation kinetics can be reproduced with both the feedback model and the sequential model but not with the two-state model. Fit parameters are given in Supplementary Table S4. (**B**) Influence of recruitment speed on the resulting activation kinetics. The sequential model was used as a proxy for the experimental data. Slow recruitment in combination with moderately fast RNA detection would yield data points that could also be fitted with the two-state model (right plot panel with error bars reflecting the experimentally measured ones). (**C**) Model for the biphasic transcriptional activation kinetics observed here using BLInCR.

To assess the influence of the recruitment speed of the activator on the observed activation kinetics we conducted simulations for different recruitment rates (Fig. 4B). To this end we used the sequential model that serves as a proxy for the experimental data as shown above. Slow recruitment of the activator (*k* ≤ 0.1 min^−1^, black/blue lines in Fig. 4B) would mask the early activation phase. In contrast, fast recruitment (*k* > 1 min^−1^, red/yellow lines) with rates similar to those achieved by BLInCR (*k* > 10 min^−1^ and *τ*_1/2_ ∼ 12 s, Table S2) could resolve both activation phases. Accordingly, the early activation phase might not be visible when using tamoxifen (*k* ∼ 0.04 min^−1^, *τ*_1/2_ ∼ 17 min (Rafalska-Metcalf et al., 2010)) or doxycycline (*k* ∼ 0.4 min^−1^, *τ*_1/2_ ∼ 100 s, (Normanno et al., 2015)) to recruit transcriptional activators. In particular, slow recruitment in combination with moderately fast RNA detection (every 5 minutes) would yield data points that could also be fitted with the two-state model (Fig. 4B, right). Thus, both fast recruitment of the activator and fast readout of RNA production are critical to resolve the biphasic activation kinetics and thereby distinguish between different activation models.

## Discussion

The BLInCR method presented here can rapidly and precisely trigger light-induced recruitment of transcription regulators to genomic loci in single living cells to measure the RNA output with high temporal resolution. The approach can be easily adapted to study the activities of other PHR-fused effectors at any nuclear subcompartment that can be marked by CIBN-fused targeting factors. While our present study applied only confocal fluorescence microscopy imaging, BLInCR is compatible with several fluorescence microscopy-based techniques like super-resolution microscopy, single particle tracking and other approaches for mobility imaging. We focused here on the measurement of RNA production via fluorescent MS2 coat proteins as a prototypical functional readout to monitor cellular processes in real-time. Further, we exploited the high temporal resolution of BLInCR to distinguish between conceptionally different models for transcriptional activation. Our kinetic analysis revealed that even strong transcriptional activators like VP16 cannot readily activate an entire gene cluster in one step (Fig. 4C). Rather, full activity is only reached after a pronounced maturation phase, which would allow actively transcribed genes to contact and activate promoters in close spatial proximity (Cremer et al., 2015; Papantonis and Cook, 2013). Notably, a similar mechanism has been proposed to explain the function of actively transcribed enhancers (Li et al., 2016). Furthermore, we illustrated that the lifetime of the activated state can be derived from experiments that apply pulsed activation patterns. We anticipate that BLInCR will facilitate the study of the dynamic regulation of transcription and other nuclear processes for which the induction kinetics as well as the persistence of the output signal are crucial parameters to understand the underlying molecular mechanisms.

## Material and Methods

### Plasmids and cell line

Effector and localizer plasmids were constructed based on sequences coding for the PHR domain of cryptochrome 2 and the CIBN domain of CIB1. pCRY2PHR-mCherryN1 and pCIBN(deltaNLS)-pmGFP described in (Kennedy et al., 2010) were obtained from Addgene (plasmids #26866 and #26867). LacI and TetR constructs are based on the fluorescently tagged proteins described in (Lau et al., 2003; Pankert et al., 2017). Human GCN5 was cloned from pAdEasy Flag GCN5 (Lerin et al., 2006) (Addgene plasmid #14106). The near-infrared fluorescent protein iRFP713 was from piRFP (Filonov et al., 2011) (Addgene plasmid #31857). The nuclear localization signal (NLS) used in some of the BLInCR constructs (Supplementary Table S1) is the peptide PKKKRKV from SV40 large T antigen (Kalderon et al., 1984) and was inserted by PCR. Other human plasmids were derived from previously described vectors containing sequences coding for TRF1 (Chung et al., 2011), TRF2 (Jegou et al., 2009), PMLIII (Jegou et al., 2009), NCL (Caudron-Herger et al., 2015), LaminB1 (Muller-Ott et al., 2014) and the viral transactivator VP16 (Gunther et al., 2013). The U2OS 2-6-3 cell line (Janicki et al., 2004) was a gift from David Spector and Susan Janicki.

### Cell culture and transfection

U2OS 2-6-3 cells were seeded in matrigel-coated (1:100 in serum-free medium for at least 30 min at room temperature) slides (Cellview, Greiner Bio-One, Austria) and transfected with the appropriate constructs using Effectene (Qiagen, Germany). Medium was changed four hours after transfection and 2 μM SAHA (Millipore) was added to the fresh medium if applicable.

### Fluorescence microscopy and image processing

Cells were kept in the dark overnight and mounted on a Leica TCS SP5 confocal microscope equipped with a HCX PL APO lambda blue 63x/1.4 NA oil immersion objective. A red flashlight was used to avoid premature exposure to blue light. The following excitation and emission wavelengths were used: CFP (405 nm/415-475 nm), GFP/YFP (488 nm/500-550 nm), tagRFP/tagRFP-T (561 nm/575-630 nm), mCherry/mKate2 (594 nm/605-750 nm), iRFP713 (633 nm/645-780 nm). Images were acquired as described below for the different experiments. Image analysis was done using the Fiji distribution (Schindelin et al., 2012) of ImageJ (Schneider et al., 2012). The enrichment *E*(*t*) of a given protein at the *tet*O or *lac*O arrays was calculated from single images or the maximum intensity projections of image stacks. To quantify enrichments, the intensity difference between the array region (*I*_array_) and the nuclear reference region (*I*_nuc_) was computed for each time point:

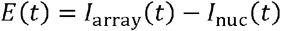

The nuclear reference region was selected to be close to the array region to account for uneven illumination of the cell if necessary, but outside of nucleoli, which generally showed some depletion of the constructs. To correct for bleaching, the decay of the mean intensity difference at the nuclear reference region and a background region outside of the cell *I*_nuc_(*t*) − *I*_background_(*t*) was fitted with a single exponential term *a*·*e*^-*k*·*t*^. The enrichment *E* at the array was then calculated as the intensity difference between array and nuclear reference region divided by the bleach contribution *e*^-*k*·*t*^, which was assumed to be the same for the array and the nucleus:

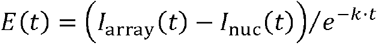

### BLInCR recruitment kinetics

U2OS 2-6-3 cells were transfected with CIBN-TetR-tagRFP-T and a PHR-YFP construct (i.e. PHR-YFP-VP16, PHR-YFP-NLS or PHR-YFP-hGCN5). Transfected cells and reporter arrays were identified in the tagRFP-T channel. Subsequently, an image series of 400 frames with 256 x 256 px images was recorded in the YFP channel at a scan speed of 1400 Hz corresponding to 204.3 ms per image and a total of ∼81 s. Excitation of YFP caused optogenetic switching, resulting in accumulation of the respective PHR-YFP construct at the CIBN site, i.e. the array seen in the tagRFP-T channel (Supplementary Fig. S2A). To quantitatively analyze the enrichment of PHR-YFP constructs at the array site, a maximum intensity projection was used to select the array region (diameter: ∼15 px) and a nuclear reference region (diameter: 30 px) at which the mean fluorescence intensities were measured (Supplementary Fig. S2A) to compute *E*(*t*) as described above. For PHR-YFP accumulation, all curves could be fitted well with a model containing two exponentials:

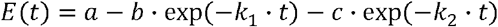

This equation describes a reaction model with two parallel first-order reactions to the same product (Steinfeld et al., 1989). From the fit, the characteristic recruitment time *τ*_1/2_ was calculated as given below:

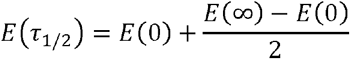

*E*(0) and *E*(∞) where calculated from the fit and the plateau value *E*(∞) = *a* was used for normalization according to *E*_norm_(*t*) = *E*(*t*)/*a*. Finally, all normalized curves were averaged, yielding curves as in Fig. 2D and Supplementary Fig. S2B. Cells that moved in *z*-direction leading to fluctuations in the enrichment curves as well as cells with very low YFP signal and bad signal-to-noise ratio were excluded from the analysis.

### BLInCR dissociation kinetics

U2OS 2-6-3 cells were transfected with CIBN-TetR, GFP-LacI and a PHR-mCherry construct, i.e. PHR-mCherry-VP16 or PHR-mCherry-NLS. Transfected cells were identified in the mCherry channel and a pre-recruitment stack (seven slices with Δ*z* = 0.5 μm, 2x line average, 512 x 512 px, 400 Hz scan speed) was recorded. To recruit the PHR-mCherry constructs to the array, two stacks were recorded in the GFP channel corresponding to 38 s illumination with blue light. The first post recruitment stack was recorded in the mCherry channel immediately afterwards and constitutes the time point *t* = 0 s. Subsequent stacks were acquired every ∼30 s for the first ∼5 min and then at longer intervals for about 30 min (Supplementary Fig. S2C). The focus was readjusted if necessary.

To quantify the reversibility kinetics, the *z*-stacks for each time point were registered using the StackReg plugin (Thevenaz et al., 1998). For each time point, a maximum projection of the registered *z* slices was made, resulting in a time series of maximum projections. Generally, cell shapes changed considerably over the 30 min acquisition, preventing registration of the time series. Consequently, the array region (diameter: 20-40 px) and a nuclear reference region (diameter: 60 px) were manually selected for each time point and mean fluorescence intensities were measured (Supplementary Fig. S2C). Importantly, the sizes of the regions were kept constant over all images of one time series.

All *E(t*) curves could be fitted well with a single exponential with a time-dependent (i.e. concentration-dependent) reaction rate similar to the model proposed by Sing *et al.* (Sing et al., 2014):

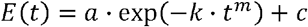

From the fit, the characteristic half-life *t*_1/2_ was calculated:

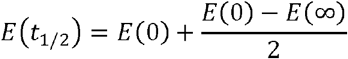

*E*(0) and *E*(∞) where calculated from the fit and used for normalization:

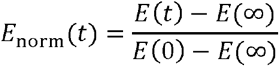

The normalized curves were averaged to yield the data shown in Fig 2E and Supplementary Fig. S2D.

### Light-induced transcription activation

U2OS 2-6-3 cells were transfected with mKate2-MS2 for RNA readout, CIBN-TetR as recruitment platform, CFP-LacI as array marker and PHR-YFP-VP16 for recruitment to CIBN-TetR and subsequent transcriptional activation. Image acquisition was similar to that used for the reversibility kinetics as described (Supplementary Fig. S2C), except for an additional third scan (CFP) and a 4x line average instead of 2x. After constitutive recruitment with a blue LED overnight, 4 slices (Δ*z* = 0.5 μm) were recorded. For the time series, the number of slices recorded was between 5 and 8 (Δ*z* = 0.5 μm) depending on cell size and shape. This was done to assure that the array was within the range recorded since it cannot be seen in the pre-recruitment image. For the first stack (pre/recruit), the channels were switched between stacks with the red channel (mKate2) being recorded first (yielding the reference image for mKate2-MS2 before VP16 recruitment), then the YFP channel and last the CFP channel. For all subsequent stacks (one stack every 2-4 min) as well as after constitutive activation, all three channels were recorded sequentially using the “between lines” mode. Hence, different color images did not need to be registered with respect to one another. However, different *z* slices had to be registered since the recording of an entire image stack with three colors lasted longer than one minute, so that movement of the cell was occasionally observed. The different *z* slices were transformed to RGB Stacks and then registered to one of the central slices based on the YFP channel using the TurboReg (Thevenaz et al., 1998) plugin.

To quantify the RNA amount at the array, maximum intensity projections were made for each time point resulting in a stack of three maximum intensity projections (mKate2, YFP and CFP). The quantification was also done manually for each time point as described above for the reversibility kinetics. Note that the array area was selected in the YFP and/or CFP channel and was kept at the same size (diameter: 20-40 px) across all time points for a single cell. The *E(t*) curves for RNA/mKate2-MS2 enrichment at the array were calculated as described above, normalized to the enrichment values before and after 50 min of VP16 recruitment, and averaged (Fig. 3B). The average curve was fitted using a simple two-state model, a model assuming positive feedback or a sequential activation model (Fig. 4A) as described below.

To calculate the RNA enrichment values at the first plateau (Fig. 3C), the inflection points were calculated from the fit of the average curves (feedback model) of untreated and treated cells. The time course of array sizes was determined from the PHR-YFP-VP16 or the CFP-LacI signal with ImageJ. The local area around the array (diameter: 60 px/3.2 μm) was selected and converted to a binary image of the array using Otsu’s method (Otsu, 1979) for thresholding. The measured array sizes at each time point were normalized to the array size at *t* = 2.5 min to assure that PHR-YFP-VP16 was fully recruited.

To compare RNA levels after constitutive activation, cells were transfected with PHR-YFP or PHR-YFP-VP16 and CFP-LacI as well as CIBN-TetR, exposed to a blue LED overnight or via expression of a co-transfected GFP-TetR-VP16 fusion protein (Supplementary Fig. S1). CLSM imaging was conducted with the same laser intensities on the same day. The fluorescence intensity enrichment of mKate2-MS2 tagged RNA was calculated from the background corrected intensities and the array area size *A*_array_ according to

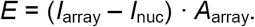

### BLInCR transcription activation and reversibility

To test the reversibility of transcription activation, cells were transfected with tagRFP-MS2 for RNA detection, CFP-LacI as array marker as well as CIBN-TetR and PHR-iRFP713-VP16 for light-induced recruitment of VP16 to the reporter array. The imaging parameters size, speed and line averaging were the same as for the light-induced transcription activation described above. Prior to recruitment and activation, a two-color stack (iRFP713 and tagRFP, sequential scan in “between lines” mode) was recorded, ensuring that neither VP16 nor RNA were detectable at the array. For light-induced recruitment of VP16, two three-color stacks (iRFP713, tagRFP and CFP, sequential scan in “between lines” mode) were recorded and the beginning of the first stack is the time point *t* = 0 s. Each stack acquisition exposed the cells to blue light for 78 s, and both stacks were recorded ∼3 min apart. Subsequently, two-color stacks (iRFP713 and tagRFP) were recorded every 2-4 min to monitor PHR-iRFP713-VP16 dissociation and tagRFP-MS2 accumulation and dissociation from the array. After 30-40 min, PHR-iRFP713-VP16 was recruited again by switching back to three-color imaging (iRFP713, tagRFP and CFP). RNA and PHR-iRFP713-VP16 quantification at the array and bleach correction was done as described above. The maximum enrichment value and the enrichment value before VP16 recruitment *E*(0) were used for normalization.

### RNA fluorescence *in situ* hybridization (FISH) and immunofluorescence (IF)

RNA production from the reporter array was analyzed by RNA FISH. Cells were seeded on cover slips and transfected with CIBN-TetR, CFP-LacI and PHR-YFP or PHR-YFP-VP16 as described above. After illumination with a blue LED overnight, cells were permeabilized on ice with CSK buffer (100 mM NaCl, 300 mM sucrose, 3 mM MgCl_2_, 10 mM PIPES, 0.5% Triton X100) containing 10 mM vanadyl ribonucleoside complex (VRC, New England Biolabs) for 5 minutes. Further processing was done at room temperature unless noted otherwise: Cells were washed once with PBS, fixed with paraformaldehyde (12 min) and washed again with PBS. Subsequently, they were incubated with 70%, 85% and 100% ethanol (3 min each) and air-dried. For MS2 stem loop RNA detection, 50 ng of the 5’-Atto-565 labeled antisense probe 5’-GTC GAC CTG CAG ACA TGG GTG ATC CTC ATG TTT TCT AGG CAA TTA-3’ (Goodier et al., 2010) per slide were mixed with 10 μg salmon sperm DNA and 5 μl formamide. The mixture was heated to 37°C for 10 min and 74°C for 7 min before 5 μl hybridization buffer (0.6 M NaCl and 60 mM trisodium citrate, pH 7.0, 20% dextran sulfate and 2 mg/ml BSA) and 10 mM VRC was added to a total volume of 11 μl. After hybridization overnight at 37°C cover slips were washed as follows: Twice with 2x SSC (0.3 M NaCl, 30 mM trisodium citrate, pH 7.0) supplemented with 50% formamide (15 min), once with 0.2x SSC/0.1% Tween (10 min, 40°C), once with 2x SSC (5 min), and once with PBS. Subsequently, YFP was visualized by immunofluorescence staining with an anti-GFP antibody (Abcam ab290, lot: GR135929-1) since the fluorophore was destroyed during RNA-FISH. Cells were permeabilized with 0.1% ice-cold Triton X-100/PBS (5 min), washed once with PBS (5 min) and blocked with 10% goat serum in PBS for 30 min. The samples were incubated with the primary antibody (1:500 in 5% goat serum/PBS) for 1-2 h or overnight at 4°C. After washing three times with PBS supplemented with 0.002% NP40, samples were incubated with an Alexa488-coupled secondary anti-rabbit IgG antibody (Invitrogen, diluted 1:500 in PBS) for 45 min and washed with PBS (3 x 5 min). Lastly, the slides were rinsed with water, 75% ethanol and 100% ethanol before mounting them with prolong gold antifade mountant including DAPI (Life Technologies). For IF staining of H3K9ac (Supplementary Fig. S3A-C), cells were fixed after 19 h exposure to blue light (LED) using 4% paraformaldehyde (7 min) and washed three times with PBS. Next, cells were permeabilized and processed as described above. The primary antibody was a rabbit antibody against acetylation of H3K9 (ActiveMotif #39917, lot: 16111002) and the secondary antibody was an Alexa568-coupled anti-rabbit IgG antibody (Invitrogen). The antibodies were diluted as described above.

### Histone extraction and western blotting

U2OS 2-6-3 cells were cultured with 2 μM SAHA (Millipore, 1:1000 from 2 mM stock solution in ethanol) or 1:1000 Ethanol for 24 h. Histones were extracted from ∼1x10^6^ cells using 0.25 M HCl as described previously^13 14^, separated by electrophoresis on a precast 4-20% polyacrylamide gel (Bio-Rad) and blotted semi-dryly onto a nitrocellulose membrane. After blocking in 5% milk/1x TBS modified histones were detected with primary antibodies against H3K27ac (1:1000 in 1% BSA/1x TBS; Abcam ab4729, lot: CR238071-2), H3K9ac (1:1000 in 1% BSA/1x TBS; Active Motif 39137, lot: 09811002) and H4ac (1:2000 in 3% milk/1x TBS; Millipore #06-866, lot: DAM1416550) and HRP-linked anti-rabbit IgG (Cell Signalling, #7074S, 1:2000 in 5% milk/1x TBS). Bands were detected by chemoluminescence using clarity western ECL substrate (Bio-Rad). After stripping the membranes with stripping buffer (Carl Roth), total H3 levels were detected with an antibody against H3 (Cell Signaling #14269, 1:1000 in 1% BSA/1x TBS) and a secondary HRP-linked antibody against mouse IgG (Cell Signalling, #7076S, 1:2000 in 5% milk/1x TBS).

### Theoretical framework for kinetic analysis

#### Kinetic two-state model for transcriptional activation

Curves for the relative RNA levels in Fig. 3B were fitted with a two-state model according to the scheme depicted below the plot. The differential equations for the activated state (*A*) and the RNA level (*R*) read

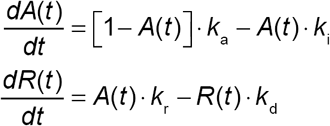

Here, *k*_a_ and *k*_i_ are the transition rates into the activated and inactive state, respectively. *k*_r_ is the RNA production rate, and *k*_d_ is the RNA dissipation rate from the array. The total number of active and inactive promoter is normalized to 1, so that the initial concentration A_0_ can take on values between 0 and 1. The solution for these equations is given by

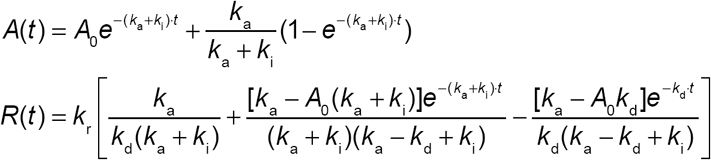

The fit parameters corresponding to the fit functions in Fig. 4A are listed in Supplementary Table S4. The decay rate *k*_d_ was set to be equal for untreated and SAHA-treated cells.

Furthermore, fits were constrained not to exceed relative RNA levels of 5 a.u. (reached in steady state at *t* = ∞), reflecting the fact that measured RNA levels remained below this level also for late time points (up to 24 hours post-induction).

#### Kinetic three-state model with feedback for transcriptional activation

Curves for the relative RNA levels in Fig. 3B were fitted with a three-state model involving feedback according to the scheme shown in below the plot. The differential equations for the activated state (*A*_1_) and the RNA level (*R*) read

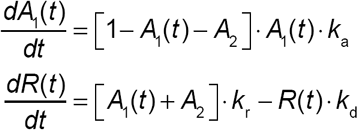

Here, *k*_a_ is the transition rate into the activated state *A*_1_, and *A*_2_ is the population residing in the activated state *A*_2_. Again, the total number of promoters is normalized to 1. The solution for these equations is given by

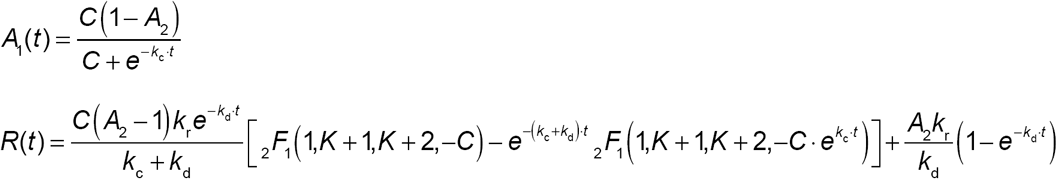

with the abbreviations 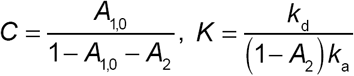 and *k*_*c*_ = (1 − *A*_2_) *k*_*a.*_

In these equations, *A*_1,0_ is the initial population in state *A*_1_, *A*_2_ is the (invariant) population in state *A*_2_, and _2_ *F*_1_*(a.b,c,x)* denotes the Gaussian hypergeometric function. The fit parameters corresponding to the fit functions shown in Fig. 4A are listed in Supplementary Table S4. The dissipation rate *k*_d_ was set to be equal for untreated and SAHA-treated cells.

#### Sequential model for transcriptional activation

Curves for the relative RNA levels in Fig. 3B were fitted with a sequential model involving *n* sequential reaction steps to transition from the inactive state (*I = S*_0_) to the active state (*A*) according to the scheme *I* → *S*_1_ → *S*_2_ →…→ *S*_*n*-1_ → *A*. Here, *S*_i_ represent intermediate states, in which no RNA is produced. For simplicity, the same rate constant *k*_a_ was chosen for each transition. The differential equations for the individual states (*I*, *S*_i_, *A*) and the RNA level (*R*) read

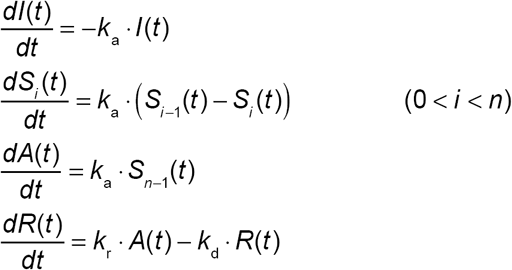

The solutions for *A*(0) = *A*_0_ and *I*(0) = 1 − *A*_0_ are given by

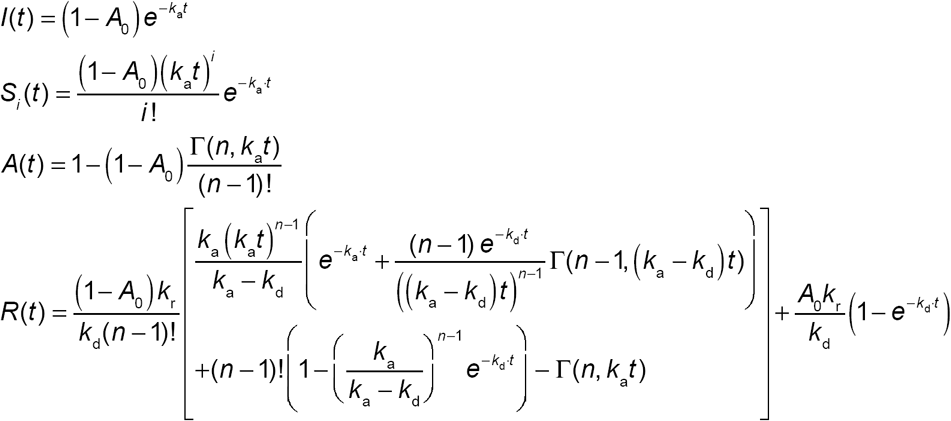

Here, *A*_*0*_ is the initial population in the active state *A*, and Γ(*x*,*y*) is the (upper) incomplete Gamma function. The fit parameters corresponding to the fit functions in Fig. 4A are listed in Supplementary Table S4. The decay rate *k*_d_ was set to be equal for untreated and SAHA-treated cells.

#### Sequential model with additional recruitment step

The scheme for the sequential activation model above was extended to explicitly include an additional recruitment step for the transcriptional activator with rate *k*_rec_. In particular, an unbound inactive state (*U*_I_) and an unbound active state (*U*_A_) were considered, with transitions from *U*_I_ to *I* and *U*_A_ to *A* that occur with rate *k*_rec_. Thus, the reaction schemes read 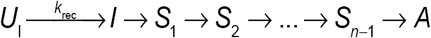 and 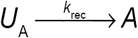. In this case, the differential equations for the individual states (*U*_A_*, U*_I_, *I*, *S*_i_, *A*) and the RNA level (*R*) read

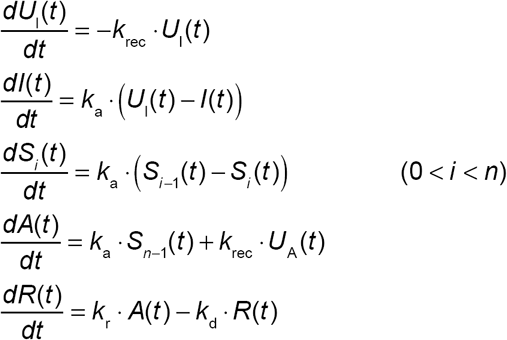

The solutions for *U*_A_ (0) = *A*_0_ and *U*_I_(0) = 1 − *A*_0_ are given by

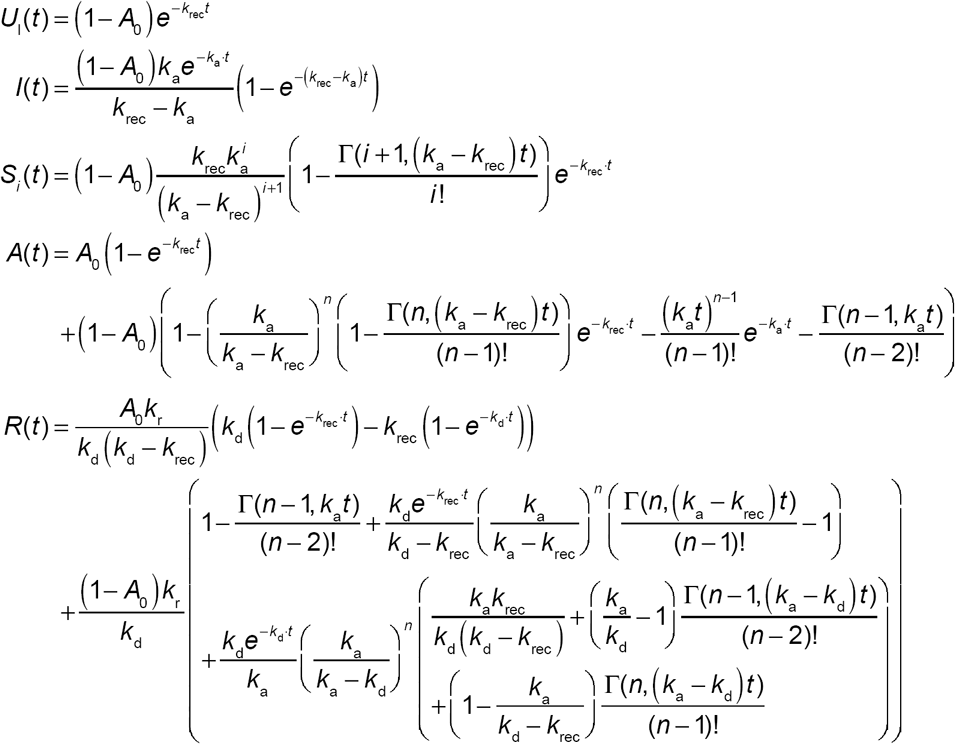

Here, Γ(*x*,*y*) is the (upper) incomplete Gamma function. Resulting curves for different recruitment rates *k*_rec_ are shown in Fig. 4B.

## Acknowledgements

We thank Caroline Bauer and Sabrina Schumacher for valuable assistance, the DKFZ Light Microscopy Facility for technical support and expertise, and Katharina Müller-Ott for comments on the manuscript. We are grateful to David Spector and Susan Janicki for providing the U2OS 2-6-3 cell line.

## Supplementary Figures

**Movie 1 | Optogenetic recruitment of PHR-YFP-VP16 to tethered CIBN-TetR.** Note that the brightness has been adjusted non-linearly (gamma=0.65) for better visibility of the signals at the array and within the nucleoplasm. Speed: 10 fps (∼2x). Scale bar: 5 μm.

**Supplementary Figure S1.**
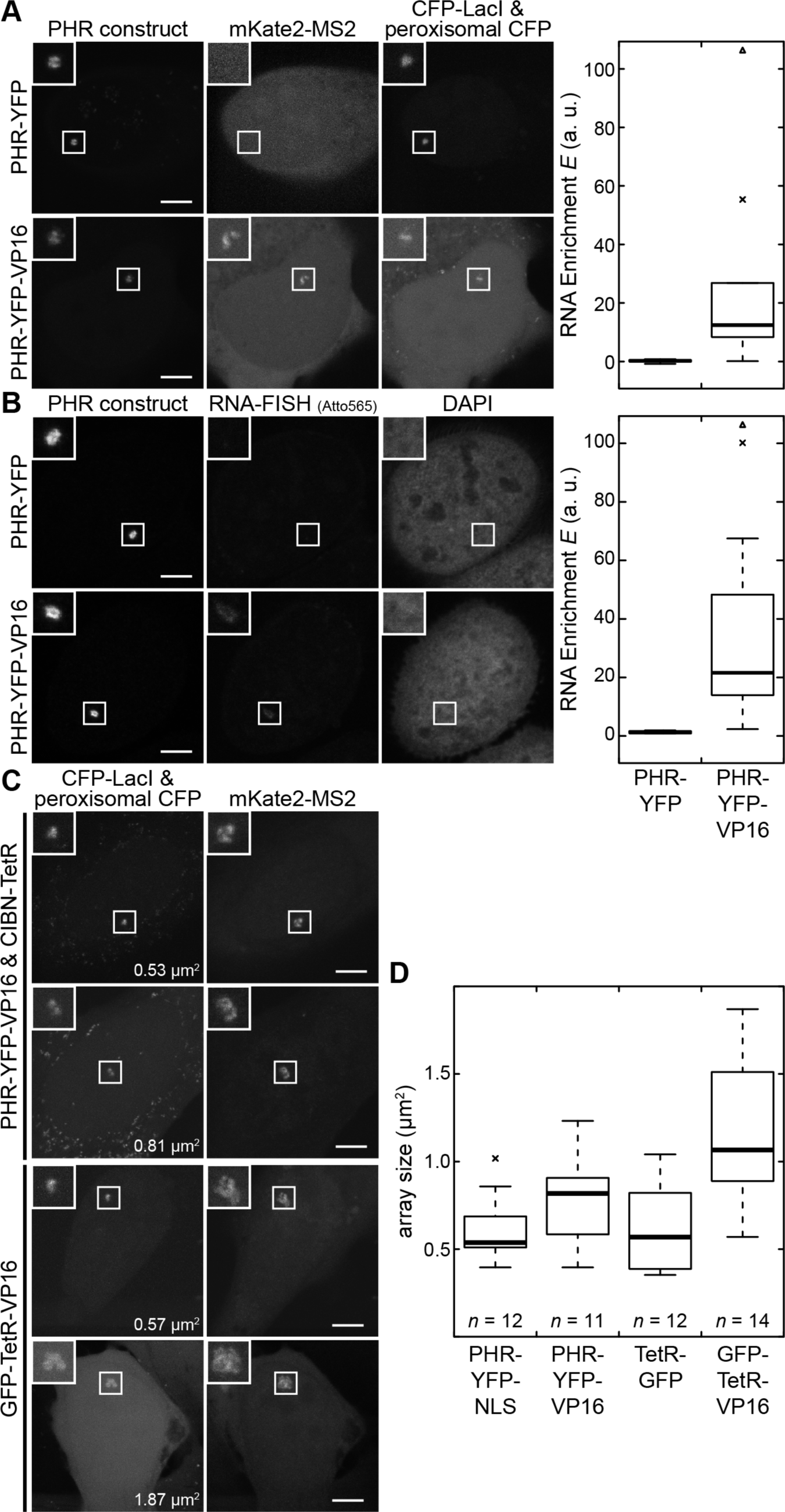
Constitutive optogenetic recruitment of PHR-YFP with or without VP16 to *tet*O repeats. Maximum intensity projection CLSM images of the indicated constructs (left) and the quantification (right) of RNA enrichment (**A & B**) or array size (**C & D**) are shown. Cells were continuously illuminated overnight with a blue LED, leading to recruitment of PHR-YFP constructs (via CIBN-TetR) to the array. (**A**) RNA was visualized with fluorescently tagged MS2 coat protein (*n*=9 for each condition). (**B**) RNA was detected with a fluorescently labeled FISH probe against MS2 loops (*n*=6 and *n*=11 for PHR-YFP and PHR-YFP-VP16, respectively). The amount of RNA detected at the array after VP16 recruitment showed a similar distribution for different RNA detection methods and varied considerably across different cells. (**C**) CFP-LacI was used for independent size measurements. All cells transfected with a VP16 construct showed mKate2-MS2 enrichment, confirming that these cells were transcriptionally active. (**D**) Quantification of array sizes with enriched VP16 compared to negative controls without VP16. On average, cells expressing TetR-VP16 fusion constructs had somewhat larger arrays compared to cells with constitutive optogenetic recruitment of PHR-YFP-VP16. Notably, some of the activated cells in the latter population had condensed arrays that were comparable to those in the negative control PHR-YFP-NLS. Scale bars: 5 μm.

**Supplementary Figure S2.**
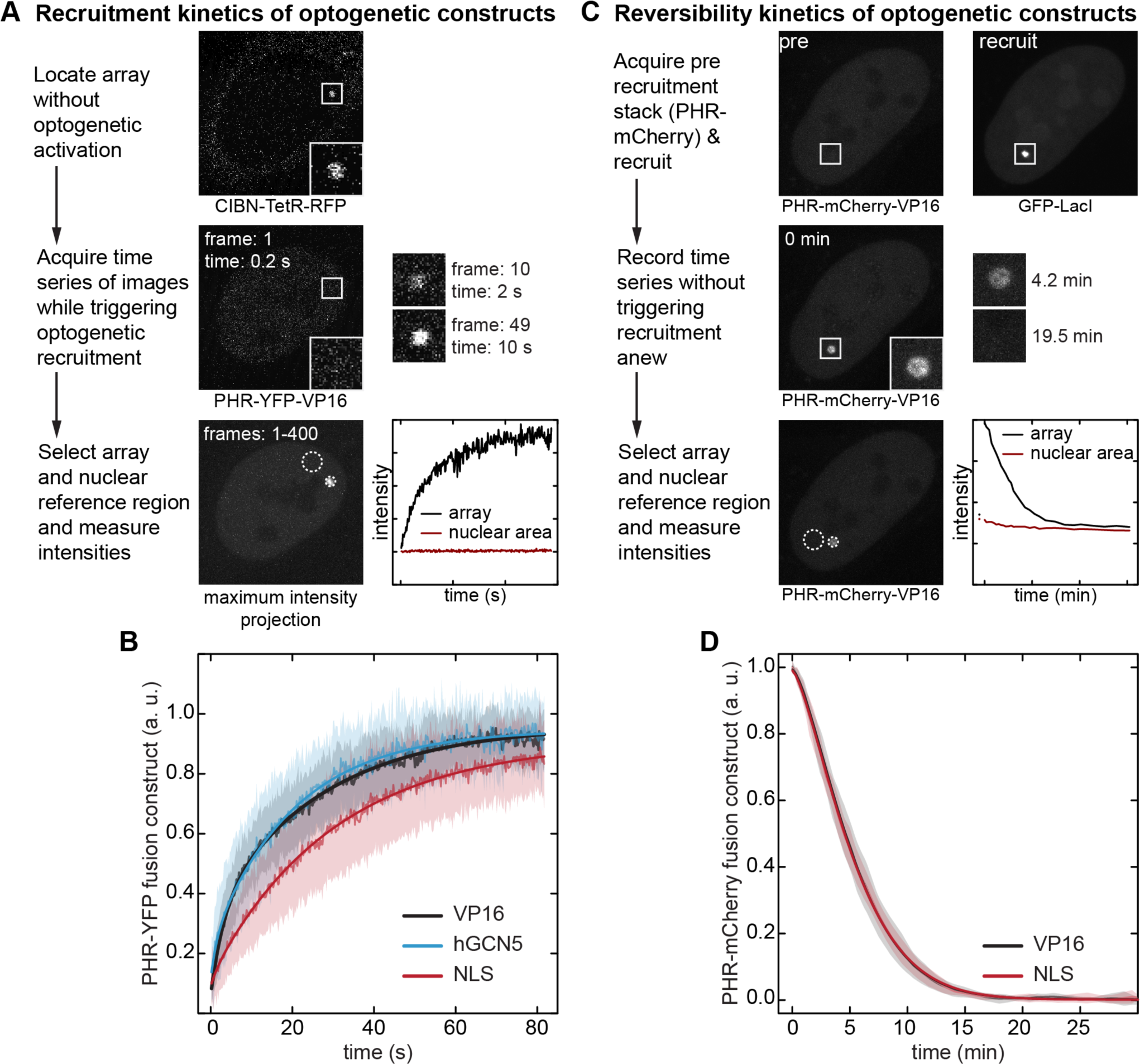
Recruitment and reversibility with BLInCR. (**A**) The typical workflow of a BLInCR experiment is illustrated for cells transfected with CIBN-TetR-tagRFP-T (localizer) and PHR-YFP-VP16 (effector). First, the cellular structure bound by the localizer (here: *tet*O array) was located in the tagRFP-T channel (excitation with yellow-green light), which did not trigger optogenetic recruitment. Next, a YFP time series was recorded while triggering recruitment of the effector to the localizer (excitation with blue light). Finally, the structure that was targeted by the localizer (here: *tet*O array) and a nuclear reference region were selected in the maximum intensity projection of the time series and the mean intensities at those regions were measured for each image. (**B**) Recruitment kinetics of PHR-YFP-fused effector proteins. Experimental means (solid lines, light), standard deviations (shaded areas) and fits of the average curves (solid lines) are shown. The fit parameters are listed in Supplementary Table S2. (**C**) A typical workflow for a BLInCR reversibility experiment is illustrated for cells transfected with CIBN-TetR (localizer), GFP-LacI (marker) and PHR-mCherry-VP16 (effector). A stack of images was recorded in the mCherry channel (excitation with yellow-orange light) prior to recruitment. The array was visualized by recording two stacks in the marker channel with blue light excitation for 38 s, thereby triggering effector recruitment to the localizer. The dissociation of the effector from the targeted structure was monitored by recording image stacks every 30-120 s without triggering the PHR switch again (excitation with yellow-orange light). Finally, the targeted structure and a nuclear reference region were selected in the maximum intensity projection of the time series and the mean intensities at those regions were measured for each image. (**D**) Plot of experimental means (solid lines, light), standard deviations (shaded areas) and fits of the average curves (solid lines). Averages of fit parameters obtained from fitting the single curves are listed in Supplementary Table S3.

**Supplementary Figure S3.**
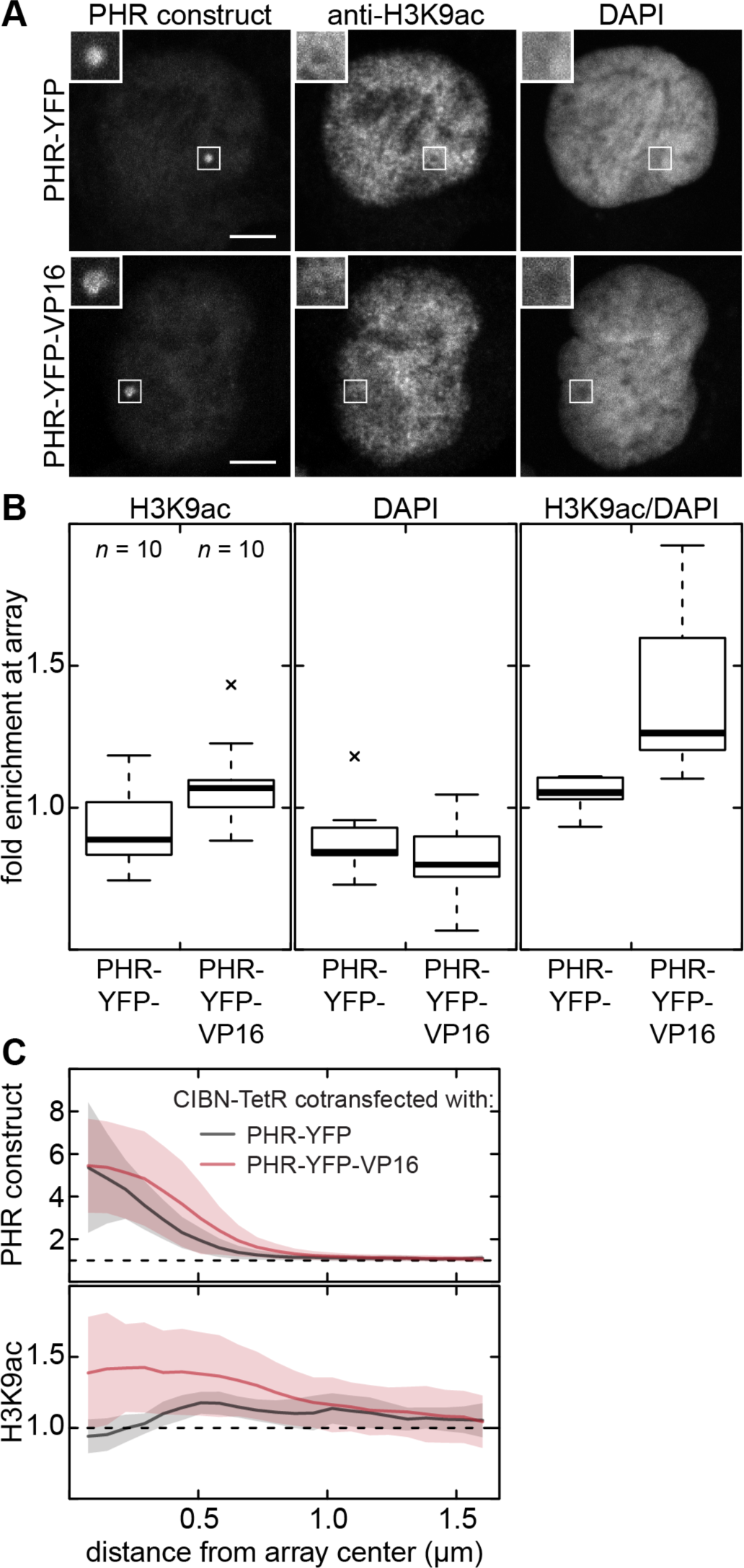
Local histone acetylation levels. (**A**) Cells were transfected with CIBN-TetR and PHR-YFP without (top) and with VP16 (bottom). When VP16 was absent, there was a drop in H3K9ac levels compared to the surrounding chromatin, which was not present when VP16 was recruited. Conversely, there was a drop in DNA density when VP16 was present, possibly indicating array decondensation (see Supplementary Fig. S1C). (**B**) Fold enrichment of H3K9ac and DNA density (DAPI) at the array compared to a nuclear reference region. The DAPI-normalized H3K9ac fold enrichment is depicted on the right. It increased when VP16 was present, indicating that the array became acetylated when activated. (**C**) Average radial profile of DAPI-normalized H3K9ac (bottom) confirmed the depletion of H3K9ac in non-activated cells and the slight enrichment in activated ones. The array center is at distance zero and the profiles of the recruited constructs are depicted for clarity (top). The averages for ten cells for each condition are shown in (**B & C**).

## Supplementary Tables

**Supplementary Table S1.** List of BLInCR constructs. For developing BLInCR, constructs with different autofluorescent protein domain fusions were constructed and tested. The constructs used for the experiments described in the main and supplementary figures are indicated in bold. All plasmid vectors will me made available via Addgene.

| CIBN constructs | PHR constructs | Others |
| --- | --- | --- |
| **CIBN-tagBFP-TetR** | **PHR-mCherry** | tagBFP-LacI |
| CIBN-YFP-TetR | PHR-YFP | **CFP-LacI** |
| **CIBN-tagBFP-hLaminB1** | **PHR-mCherry-NLS** | **GFP-LacI** |
| CIBN-YFP-hLaminB1 | **PHR-YFP-NLS** | RFP-LacI |
| CIBN-tagBFP-hTRF1 | **PHR-mCherry-VP16** | **TetR-GFP** |
| CIBN-YFP-hTRF1 | **PHR-YFP-VP16** | TetR-YFP |
| **CIBN-tagBFP-hTRF2** | **PHR-iRFP713-VP16** | TetR-mRFP |
| CIBN-YFP-hTRF2 | PHR-mCherry-hPMLIII | GFP-MS2 |
| **CIBN-tagBFP-hNCL** | PHR-YFP-hPMLIII | **tagRFP-MS2** |
| CIBN-YFP-hNCL | **PHR-YFP-hGCN5** | **mKate2-MS2** |
| **CIBN-tagBFP-hPMLIII** | PHR-YFP-hGCN5-NLS | mCherry-MS2 |
| CIBN-YFP-hPMLIII | PHR-YFP-hGCN5mut | tagBFP-TetR-VP16 |
| **CIBN-TetR-tagRFP-T** | PHR-YFP-hHP1β | GFP-TetR-VP16 |
| CIBN-LacI-tagRFP-T | PHR-mCherry-hHP1β | tagBFP-LacI-VP16 |
| CIBN-hTRF1-tagRFP-T | PHR-YFP-hPMLIII | GFP-LacI-VP16 |
| CIBN-hTRF2-tagRFP-T | PHR-mCherry-hPMLIII |  |
| **CIBN-TetR** |  |  |
| CIBN-LacI |  |  |
| CIBN-YFP |  |  |
| CIBN-mCherry |  |  |

**Supplementary Table S2.** Summary of kinetic parameters for optogenetic recruitment. Single recruitment curves were fitted with a model describing two parallel first-order reactions (*E*(*t*) = *a* − *b* · exp(–*k*_1_ · *t*) − *c* · exp(–*k*_2_ · *t*)). The plateau value *a* was used to normalize each curve. The resulting fit parameters of averaged curves with their standard fit error are listed. ^a^ The characteristic recruitment time *τ*_1/2_ for reaching half maximal levels was computed as average and standard deviation of all single values.

|  | PHR-YFP-VP16 | PHR-YFP-hGCN5 | PHR-YFP-NLS |
| --- | --- | --- | --- |
| number of cells $n$ | 20 | 19 | 20 |
| $k_1$ (s<sup>-1</sup>), fast reaction | 0.31 ± 0.02 | 0.67 ± 0.19 | 0.62 ± 0.46 |
| $b$, fraction fast | 0.27 ± 0.01 | 0.12 ± 0.02 | 0.03 ± 0.01 |
| $k_2$ (s<sup>-1</sup>), slow reaction | 0.037 ± 0.001 | 0.049 ± 0.001 | 0.033 ± 0.001 |
| $c$, fraction slow | 0.63 ± 0.01 | 0.71 ± 0.01 | 0.79 ± 0.004 |
| $\tau_{1/2}$ (s)<sup>a</sup> | 11.7 ± 5.5 | 13.2 ± 7.0 | 25.8 ± 12.0 |

**Supplementary Table S3.** Summary of kinetic parameters for dissociation after optogenetic recruitment. Single reversibility curves were fitted with an exponential model comprising a time-dependent reaction rate (*E*(*t*)= *a* · exp (−*k* · *t*^*m*^ + *c*). Mean and standard deviation are shown.

|  | PHR-mCherry-VP16 | PHR-mCherry-NLS |
| --- | --- | --- |
| number of cells $n$ | 13 | 12 |
| $k$ (s<sup>-1</sup>) | 0.08 ± 0.04 | 0.09 ± 0.03 |
| $m$ | 1.44 ± 0.17 | 1.39 ± 0.19 |
| $t_{1/2}$ (min) | 4.6 ± 0.7 | 4.5 ± 0.5 |

**Supplementary Table S4.**
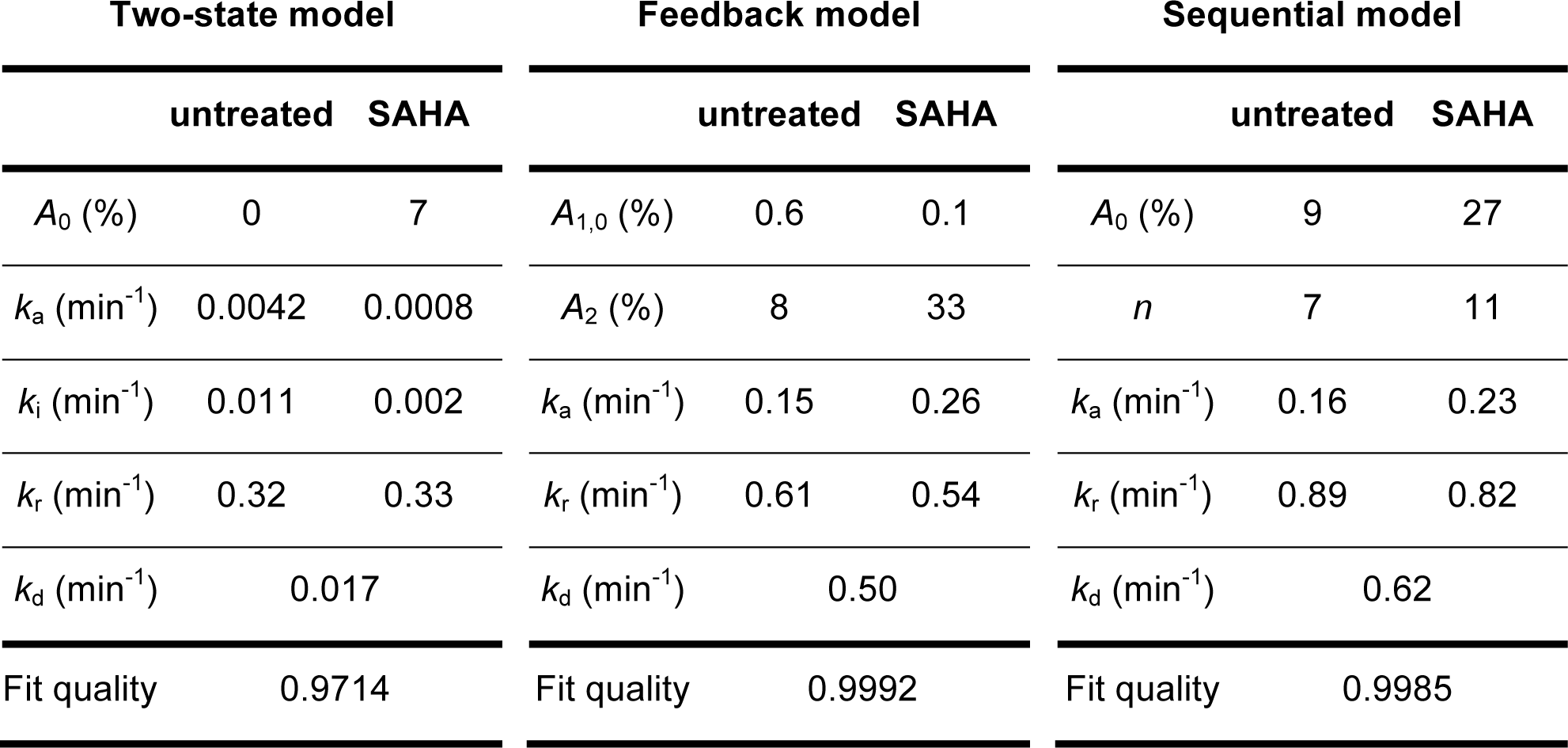
Fit parameters for two-state, feedback and sequential model fit of BLInCR-induced transcription activation. Differential equations and their solution are described in the Methods section. Averaged curves and fit curves are depicted in Fig. 3B. The fit quality corresponds to the adjusted coefficient of determination *R*^2^.

